# Breeding at higher latitude as measured by stable isotope is associated with higher photoperiod threshold and delayed reproductive development in a songbird

**DOI:** 10.1101/789008

**Authors:** D. Singh, S.R. Reed, A.A. Kimmitt, K. A. Alford, E.D. Ketterson

## Abstract

Many organisms time reproduction to photoperiod, a constant from year to year. Predicting how anthropogenic change will influence future timing demands greater knowledge of the current role of photoperiod. We held two closely related bird populations in a common environment. One population is resident; the other winters in sympatry with the resident population but migrates north prior to reproducing. We increased photoperiod gradually and measured preparation for migration and reproduction, using feather stable isotopes to estimate breeding latitude. We predicted population differences in the minimum stimulatory day length to elicit a response (CPP, critical photoperiod) and co-variation between CPP and distance migrated. We found clear population differences in CPP and greater CPP in longer distance migrants. We conclude that current geographic variation in reproductive timing has a genetic or early developmental basis and recommend that future research focus on how anthropogenic changes will interact with CPP to adjust timing of reproduction and migration.

## Introduction

Animals across the globe follow the seasons and match their growth, development, gonadal recrudescence, migration, and other seasonal life-history states to exploit the seasons most favorable for survival and reproduction (Wingfield et al., 1992; Dawson 2013). Birds breeding at different latitudes vary in duration and timing of seasonal life-history states to match breeding to periods when resources well-suited for nesting growth are abundant (Lack 1968; Visser at al., 2004). Different species depend on food supplies available at different times of the year, hence optimal timing varies by species and populations within species (Dawson and Goldsmith 1983; Wingfield et al., 1992; Dawson et al., 2001; Ball and Ketterson 2008; Watts et al., 2015). When there is a mismatch between food availability and the timing of breeding, nestling growth and survival can be compromised (Visser at al., 2004; Jonzén et al., 2006).

Individuals must prepare in advance to time their seasonal events to match the environment they occupy (Menaker 1971; Bradshaw and Holzapfel 2007). Photoperiod is the only consistent reliable cue for seasonally breeding animals and is predictable at a given latitude. Hence, changing photoperiod (i.e., day length) acts as the primary predictive cue to time seasonal phenological events such as migration and breeding (Rowan, 1926; Wingfield et al., 1992; Bronson and Heideman, 1994; Dawson 2001). In general, rate of gonadal maturation appears to be directly proportional to increasing day length (Farner and Wilson, 1957; Follett and Maung, 1978). Photoperiodic responses depend on encephalic photoreceptors perceiving light during the stimulatory phase of a daily rhythm of sensitivity (Follett et al., 1992; Ball and Balthazart, 2003; Yasuo et al., 2003). Seasonally breeding animals that undergo annual gonadal recrudescence and regression in response to changing photoperiod as a primary predictive cue, also rely on supplementary cues to initiate and regulate timing of reproductive development (Bronson and Heideman, 1994; Dawson 2001; Wingfield 2012). Towards the end of the breeding season, many bird species are no longer responsive to long days and are said to become photorefractory. They show a decline in gonad volume and reduced testosterone while days are still long, well before the return of short photoperiods during autumn (Burger, 1949; Miller, 1954). Exposure to short days during autumn is then required to break the photorefractory period and restore a bird’s ability to undergo gonadal recrudescence in response to increasing photoperiod the following spring (Farner and Mewaldt, 1955). In short, seasonal phenology can be referred to in terms of the periodic appearance of life-history states consisting of a photosensitive state capable of responding to increasing photoperiod when encountered, a photostimulatory state that is induced by increasing day length, and a photorefractory state in which an animal is no longer responsive to long days.

Reproductive timing is driven by the hypothalamic-pituitary-gonadal (HPG) axis. Gonadotropin releasing hormone 1 (GnRH1) is released from the hypothalamus to stimulate release of the gonadotropins, luteinizing hormone (LH) and follicle-stimulating hormone (FSH), from the pituitary (Li et al., 1994; Cho et al., 1998). LH and FSH stimulate gonadal growth and development of gametes, as well as production and release of sex steroids. Injecting controlled doses of exogenous GnRH (i.e., a GnRH challenge) to individuals and measuring downstream activity of the HPG axis has been a successful tool to investigate variation in animals’ physiological state and behavior (Jawor et al., 2006; Spinney et al., 2006; Grieves et al., 2016).

While much of this has been known for decades, significant knowledge gaps remain with respect to the specific mechanisms accounting for timing differences among populations that breed in different environments (Fudickar et al., 2016; Ramenofsky et al., 2017). We studied dark-eyed juncos (*Junco hyemalis*), a small songbird that consists of migratory and sedentary (i.e., resident) populations, some of which live in sympatry during the winter and early spring (Fudickar et al., 2016; Grieves et al., 2016). Residents initiate preparation for reproduction prior to the departure of migrants for their breeding grounds. Following spring migration, migrant and resident juncos are geographically isolated for the remainder of the breeding season. Hence migrants and residents can be exposed to the same environment in spring but different environments during summer.

In a prior study, resident and migrant male juncos were held captive in a common garden and exposed to the same photoperiod programmed to match the natural increase in spring. Residents were found to increase cloacal protuberance volume (CPV; a primary sperm storage structure for male birds) earlier than migrants, i.e. at a shorter photoperiod (Fudickar et al., 2016). Here, we extend this study to examine changes in the reproductive axis during all four life history states and report differences in the critical photoperiodic threshold (CPP) in spring, as well as differences in the timing of breeding termination and attaining refractoriness. We predicted that migrants and residents held in a captive common environment in gradually increasing photoperiod would differ in the photoperiod at which cloacal protuberance (CPV), baseline testosterone (T_0_), and testosterone in response to GnRH challenge (dT) would increase in spring, with the CPP being lower in residents. We also predicted that residents would enter the photorefractory state later than migrants, thus prolonging the time when CPV, T_0_, dT were elevated. We used stable hydrogen isotopes ratios (δ^2^H) in feathers to estimate breeding latitude, which has been used as a proxy for determining variation in locations where feathers are grown (Rubenstein et al., 2002; Hobson 2003).

If our predictions were supported, we would conclude that population level variation in CPP and thus in timing has a genetic or early developmental basis, which would permit further investigation of the locus of variation in brain, gonad or periphery. If our predictions were not borne out, i.e., migrants and residents did not differ in CPP in a common environment, this would suggest that timing is highly flexible regardless of migratory strategy or population of origin.

## Material and Methods

### Study Species

The dark-eyed junco (*Junco hyemalis*) is a broadly distributed North American songbird (Nolan et al., 2002). Diversifying approximately 15,000 years ago, juncos subspecies vary in plumage coloration, reproductive timing, and migratory behavior (Atwell et al., 2011; Fudickar et al., 2016). Within this species complex, a migratory subspecies, *J.h. hyemalis,* (hereafter ‘migrants’) breeds in temperate coniferous and mixed forests across Canada and Alaska, whereas a sedentary subspecies, *J.h. carolinensis*, (hereafter, ‘residents’) is found year-round in Appalachian Mountains of the eastern United States. Following the fall migration, migrants overwinter in the United States east of the Rocky Mountains. Some migrant and resident subpopulations are found in overlapping distributions in the Appalachian Mountains during the winter; specifically, both migrants and residents are frequently caught foraging in mixed flocks in the winter at Mountain Lake Biological Station, in Pembroke, VA (Nolan, 2002).

### Bird Capture and Housing

Between November 1 to December 5, 2017 male overwintering migratory dark-eyed juncos (n=45) were captured using mist nests from their overwintering sites in Bloomington, IN (39.16 °N, 86.52°W). Additionally, sympatric resident (n=15) and migrants (n=15) male dark-eyed juncos were captured at University of Virginia’s Mountain Lake Biological Station in Giles County (37.37 °N, 80.52°W). Scientific collecting permits were issued by the Virginia Department of Game and Inland Fisheries (permit # 052971), the Indiana Department of Natural Resources (permit# 1803), and the US Fish and Wildlife Service (permit # 20261). All methods were approved under protocol (# 15-026-17) by the Indiana University Institutional Animal Care and Use Committee.

Resident dark-eyed juncos are relatively bigger in body size, more uniformly colored, and heavier in body mass than migrants except during migration, when migrants store fat (Pyle, 1997). We classified the subpopulations using plumage and bill coloration (pink bill, *J. h. hyemalis*; blue-gray bill, *J. h. carolinesis:* Nolan 2002). Age was determined by looking collectively at wing, plumage color, brown/black contrast in the iris of the eye and ossification of skull (Nolan 2002; Cristol et al., 2003). Sex was determined by measuring wing length and confirmed later by growth of cloacal protuberance. After capture, all the birds were transported to Kent Farm Research Station in Bloomington, Indiana and housed in outdoor aviary under natural day length, temperature and *ad libitum* food until December 15, 2017. On January 18, 2018, we moved all birds to individual cages (61 × 46 × 46 cm and 46 × 46 × 46 cm) with *ad libitum* food and water. Migrants and residents were randomly distributed across three rooms for four months. After four months the birds were free-flying until the endpoint sampling at 16L photoperiod on July 31, 2018.

### Feather Stable Hydrogen Isotopes

The most distal secondary feather of the right wing was collected from each individual at the time of capture for analysis of δ^2^H. After collection, feathers were cleaned, cut from most distal end, weighed to approximately 0.5 mg, and placed into a 3 × 5-mm silver capsule, and mailed for further quantification of δ^2^H to the US Geological Survey Stable Isotope Lab in Denver, CO. δ^2^H values were measured using established methods of mass spectrometry (Wunder et al., 2012; Fudickar et al., 2016). The δ^2^H ratios were reported in parts per mil notation (‰) with respect to VSMOW (Vienna Standard Mean Oceanic Water) using internal standards. We used North American δ^2^H precipitation map for August (http://wateriso.utah.edu/waterisotopes/index.html) for the schematic representation of junco δ^2^H values (Fig. 1 a). The δ^2^H values were used as a continuous variable against all the physiological and hormone measures. To analyze latitudinal differences in the CPP for physiological responses, we created two subjective groups within migrants for analysis: high latitude migrant (HLM; −141‰ to −90‰), low latitude migrants (LLM; −88‰ to −30‰). The values for low latitude migrants were similar to those of residents (Fig. 1 b).

**Figure 1.**
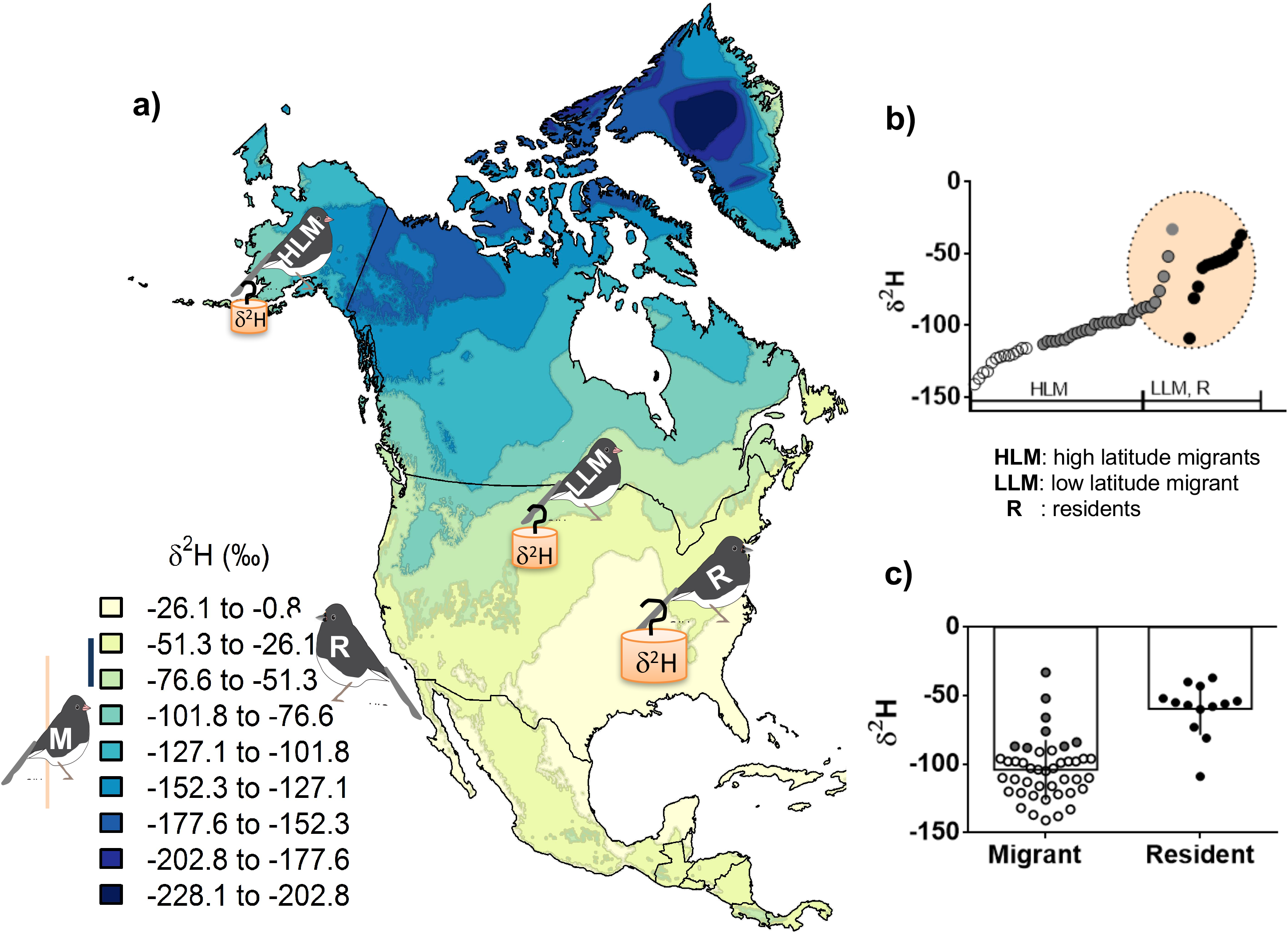
Schematic diagram of the hydrogen isotope (δ^2^H) precipitation map at different latitudes in North America during the months of July and August, Juncos undergo a pre-basic molt after breeding and their feathers incorporate stable hydrogen isotope signatures from their local environments (Rubenstein et al., 2002). Left panel represents the latitudinal distribution of dark-eyed juncos breeding range based on their feather δ^2^H (a). The bar graph represents mean difference in resident and migrant juncos’ hydrogen isotopes. Values of δ^2^H on the Y-axis are represented as hollow circle for migrants, solid grey circle for LLM, and solid circle for residents. The alpha was set at 0.05 (b). Some of the individual migrants’ overlap in isotope with residents (c). The bluish bill residents (*J.h. carolinesis*) (R; δ^2^H: −37 to −81) breed at lower latitudes and have heavier δ^2^H in comparison to pink bill migrants (*J.h. hyemalis*) which breed over a wide latitudinal range from south (low latitude migrants; δ^2^H: −30 to −88) to farther north (high latitude migrants; δ^2^H: −90 to −141). Have you double checked the numbers recently?

### Experimental Design and Sample Collection

In order to determine differences in gonadal recrudescence and migration-related physiological changes between residents and migrants in response to increasing photoperiod, we artificially regulated changes in day length. Photoperiod was increased every twelve days from January 18 to May 6 in the following schedule: 9L:15D, 10L:14D, 11L:13D, 11.4L:12.6D, 11.75L:12.25D, 12L:12D, 12.4L:11.6D, 12.75L:11.25D, 13L:11D, 15L:9D, 16L:8D. After May 6, day length remained the same until the end of the experiment on July 31, 2018. Birds were processed for physiological measurements and bled after experiencing three days in each photoperiod till 15L. Under 16L of day length, all physiological measurements except bleeding were continued until the birds regressed their CP after experiencing 40 days in 16 L. At the end of period at 16L day length, all the birds were bled to measure T_0_ and dT in photorefractory state.

### Morphological Measurements

During each sampling, we measured other indicators for preparation for reproduction and migration, including subcutaneous fat score (FS), cloacal protuberance volume (CPV) and body mass (BM) (Fudickar et al., 2016; Greives et al., 2016). Cloacal protuberance volume (CPV) is used as a measure of spermatogenesis, sperm storage and gonadal growth during the breeding season in males (Wolfson 1952). Volume of the CP was estimated using the equation for the volume of a cylinder, V=π(radius)^2^Height (Schut et al., 2012). Postnuptial (pre-basic) molt was scored at the end of the experiment based on primary, secondary, and head feathers in both the populations. Each region was given a score from 1-10 depending on the extent of molting feathers: no molt as 0 (0%), light molt (1-10%), moderate molt (11-50%) and heavy molt (51-100%). The percentages were summed to generate a total molt score for each bird (modified from Ramenofsky et al., 2017).

### Blood Sample Collection and Testosterone Hormone Assays

Immediately after capturing a bird from its cage, we took a 100 μl of blood sample by puncturing the alar wing vein for baseline testosterone (T_0_). Birds then received an intrapectoral muscle GnRH injection (~50 μl) (chicken GnRH, American peptide, Sunnyvale, CA) dissolved in PBS vehicle, which is known to activate the HPG axis in juncos (Wingfield et al., 1979; Jawor et al., 2006; Greives et al., 2016). Thirty minutes following the GnRH injection a second blood sample (50 μl) was taken from the wing vein to measure GnRH-challenged testosterone (T_30_) levels. Birds were kept in an opaque bag between injections to reduce stress. After collection, blood samples were immediately processed to extract plasma and stored at − 20°C until assayed for testosterone.

We determined T_0_ and T_30_ concentration from 20 μl plasma aliquots following established methods for our species (Jawor et al., 2006; Fudickar et al., 2016), using high sensitivity testosterone kits (Enzo Life Sciences, ADI-900-176, Farmingdale, NY) to determine circulating levels of T_0_ and T_30_. The GnRH induced testosterone level (dT) was calculated by subtracting T_0_ from T_30_. All samples were measured in duplicate and randomized over forty plates. The intra-plate and inter-plate coefficient of variation were 6.77% ± 2.07% (mean ± SE) and 13.7% respectively.

### Statistical Analysis

Data were analyzed using R (version 3.2.0). Differences in mean hydrogen isotope ratios (δ^2^H) between migrants and residents were determined using an unpaired Student’s t-test of population means. We used a Box-cox test of transformation to determine the normal distribution of all the response variables (i.e., T_0_, dT, CPV, FS, and BW). We used a square root transformation for CPV and FS, a logarithmic transformation for T_0_ and dT, and no transformation for BW. To quantify whether day length, population, or the interaction between day length and population had a significant association with response variables, we used two-way analysis of variance (2-way ANOVA) followed by Tukey’s post-hoc multiple comparison tests (alpha < 0.05). Considering repeated measures for the same individuals, we used a generalized liner mixed-effect model (GLMM) with day length and population as main effects, and age, δ^2^H as a covariate to determine effect of treatment on physiological responses. To find the critical photoperiod at which physiological parameters started to change, we used change point analysis (CPA) package in R (Killick and Eckley 2014; Robart et al., 2018). We used change point mean function which is based on the likelihood ratio and cumulative sum (CUSUM) test statistics. The CUSUM distribution does not assume data to be normally distributed and specified a single change point.

To assess co-variation between δ^2^H values as a continuous variable and morphological and hormonal measurements, we combined migrants and residents and performed Pearson correlations for CPV, BW, T_0_, dT, and molt score, and Spearman correlation for FS on one sampling date for each of four life-history states (LHSs): (1) photosensitive (9L, defined as beginning of experiment prior to recrudescence), (2) recrudescence (defined as the date of change point for CPV and dT), (3) photostimulatory, (defined as the date of seasonal peak values at 15L), and (4) photorefractory, (defined as the date of lowest seasonal value, 16L endpoint). We also calculated these correlations for migrants only on these same dates.

## Results

### Hydrogen Isotope Values for Migrants and Residents

There was a large range in the individual δ^2^H values in migrants (lowest δ^2^H = −141‰, highest δ^2^H = −33‰) in comparison to resident juncos (lowest δ^2^H = −81‰, highest δ^2^H = −37‰; except one outlier that had δ^2^H = −109‰). Mean δ^2^H differed significantly between resident and migrant juncos (p < 0.0001; Student’s t-test). Mean δ^2^H isotope was significantly lower in migrants than in residents (migrants mean δ^2^H = −104.9‰, residents mean δ^2^H = −58‰; Fig. 1 c).

### CPP for Gonadal Recrudescence in Migrants and Residents

CPV varied significantly by day length (F_13, 663.38_ = 66.3716, p < 0.0001), population (F_1, 52.34_ = 23.0848, p < 0.0001), and the interaction between day length and population (F_13, 663.43_ = 7.1168, p < 0.0001; Fig. 2 a; Table 1). The change point analysis showed CPP to be lower for gonadal recrudescence in residents than migrants. Growth in CPV in residents was detected at 12.4 h of day length, whereas migrants did not exhibit significant growth of CPV until 13 h of day length (Fig. 2 a). T_0_ also varied significantly with day length (F_13, 659.60_ = 12.417, p < 0.0001) and the interaction between day length and population (F_13, 659.64_ = 1.9205, p = 0.02521), but there was no effect of population (Fig. 2 b; Table 1). Change point analysis showed no CPP for T_0_. The variable dT varied with day length (F_13, 663.29_ = 62.786, p < 0.0001), population (F_1, 52.11_ = 45.5151, p < 0.0001), and the interaction between day length and population (F_13, 663.34_ = 5.5765, p < 0.0001; Fig. 2 c; Table 1). The effect of the co-variate δ^2^H was close to significance (F_1, 51.94_ = 3.9416, p = 0.0524). Similar to CPV response, residents showed earlier dT elevation at 11 h of day length, whereas migrants were delayed by 1h to 12 h of day length (Fig. 2 c). Comparing dT between HLM, LLM and Residents showed no difference. Interestingly, LLM elevated dT at 11.4 h of day length which differed from migrants originating from higher latitudes (Fig. 2 d). Age did not show any variation in any physiological response.

**Figure 2.**
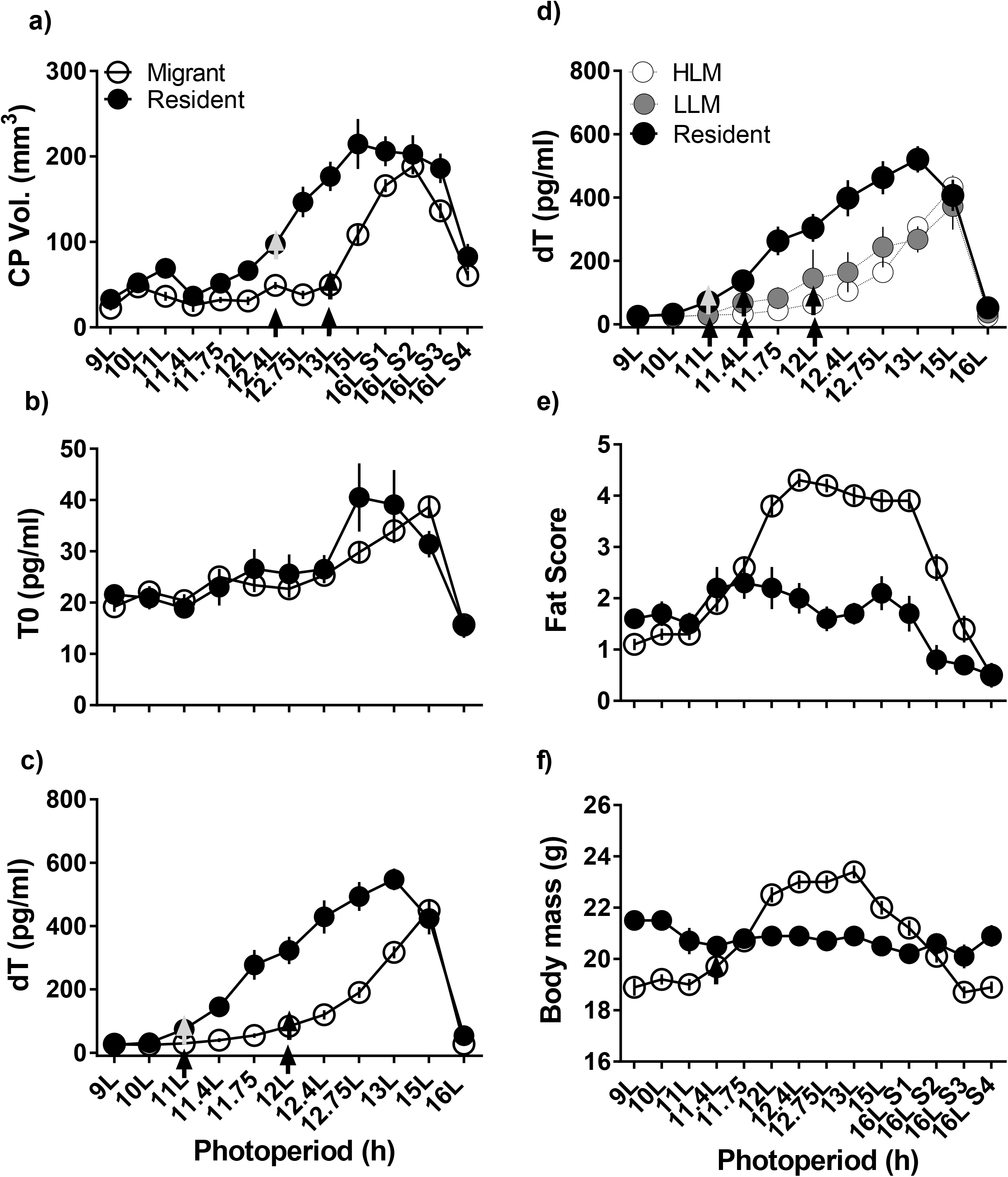
Latitudinal cline in critical photoperiod threshold for different seasonal physiological responses. Measurement of CPV (a), T_0_ (b), dT (c), FS (d), BW (e) in migrants (hollow circle) and residents (solid circle), and dT (f) in HLM (hollow circle), LLM (grey solid circle) and residents (solid circle) starting from 9 h of light to 16h of light. Y-axis represents the physiological parameters and X-axis represents increasing photoperiod in hours of day length exposure. Each data point represents mean ± SEM. Data were analyzed using a mixed-effect model with repeated measures for effects of day length, population, and interaction between day length and population. Statistical significance was defined by alpha <0.05. The arrow in the circle and above day length shows critical photoperiod threshold point in respective population, determined by change point analysis. The seasonal life-history states (LHSs) were defined as photosensitive (9L), recrudescence (CPV, 12.4L; dT, 11L), photostimulatory (15L), and photorefractory (16L) state.

**Table 1.**
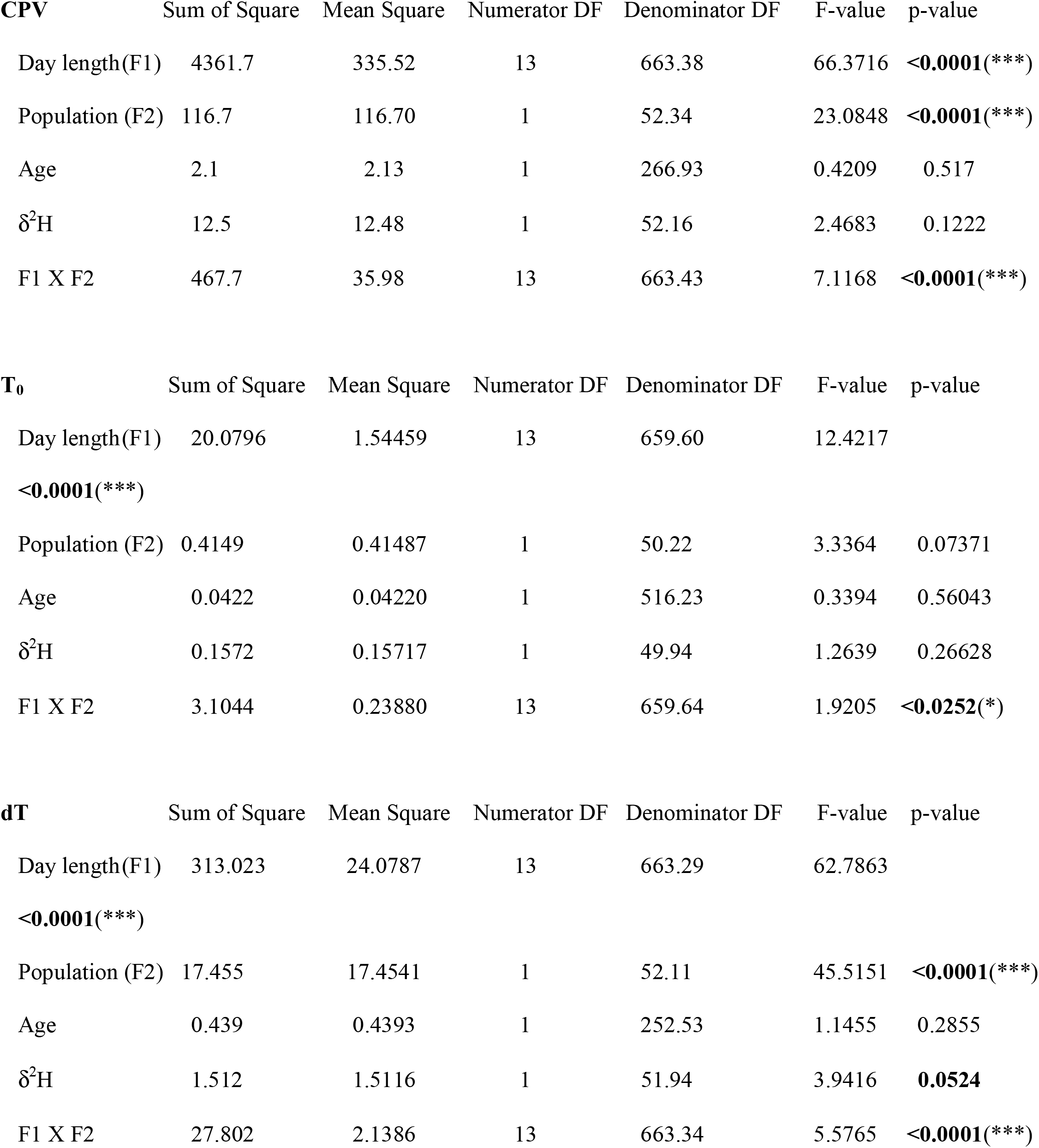
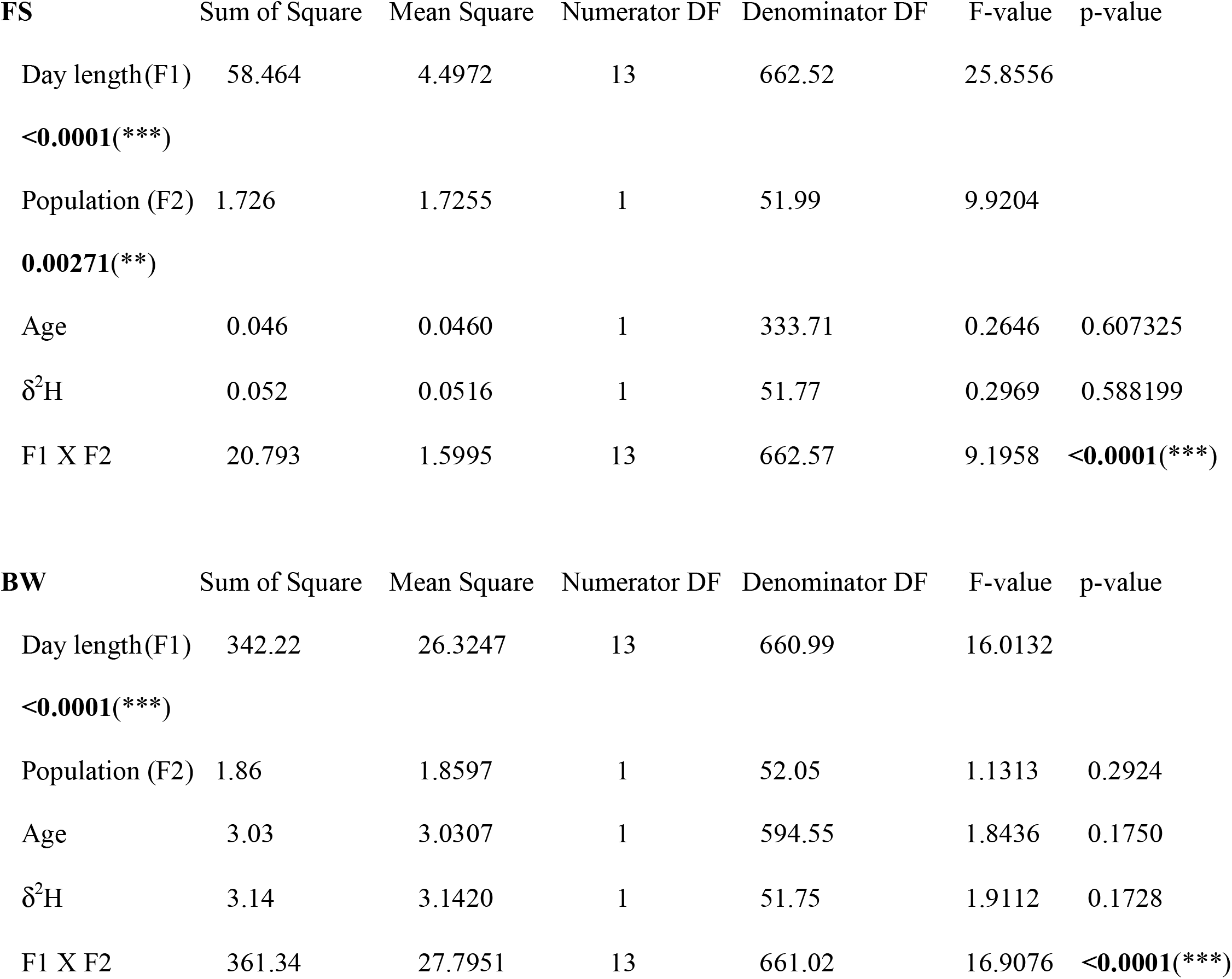
Factor (s) affecting different physiological responses (CPV, FS, BW, T_0_, dT): Analysis of variance table of type III with Satterthwaite approximation for degrees of freedom derived from linear mixed effects model. Number of asterisk (*) denotes level of significance p <0.05(*), p <0.001(**), P<0.0001(***).

### CPP for Fat score and Body mass

Migrants showed increase in pre-migratory fat score with increasing day length (F_13, 662.52_ = 25.8556, p < 0.0001) in comparison to resident birds which did not fatten (F_1, 51.99_ = 9.9204, p = 0.0027). There was also a significant interaction between day length and population (F_13, 662.57_ = 9.1958, p < 0.0001; Fig. 1 e, Table 1). At the beginning of the experiment, residents had higher body mass than migrants due to their larger body size. Migrant body mass increased significantly with day length as they fattened (F_13, 660.99_ = 16.0132, p < 0.0001), and the interaction between day length and population was significant (F_13, 661.02_ = 16.9076, p < 0.0001; Fig. 2 f, Table 1). Change point analysis revealed CPP for body mass at 11.4 h of day length for migrants (Fig. 2 f); resident birds did not change body mass as day length increased (Fig. 2 f).

### Life-history State Dependent Changes in Relationship of Phenology to Stable Isotope Values

We examined LHS-dependent changes in the relationships among CPV, dT, BW, FS with respect to δ^2^H values, considering residents and migrant collectively (Fig. 3) and dT/BW relationship with δ^2^H values in migrants separately (Fig. 3).

**Figure 3.**
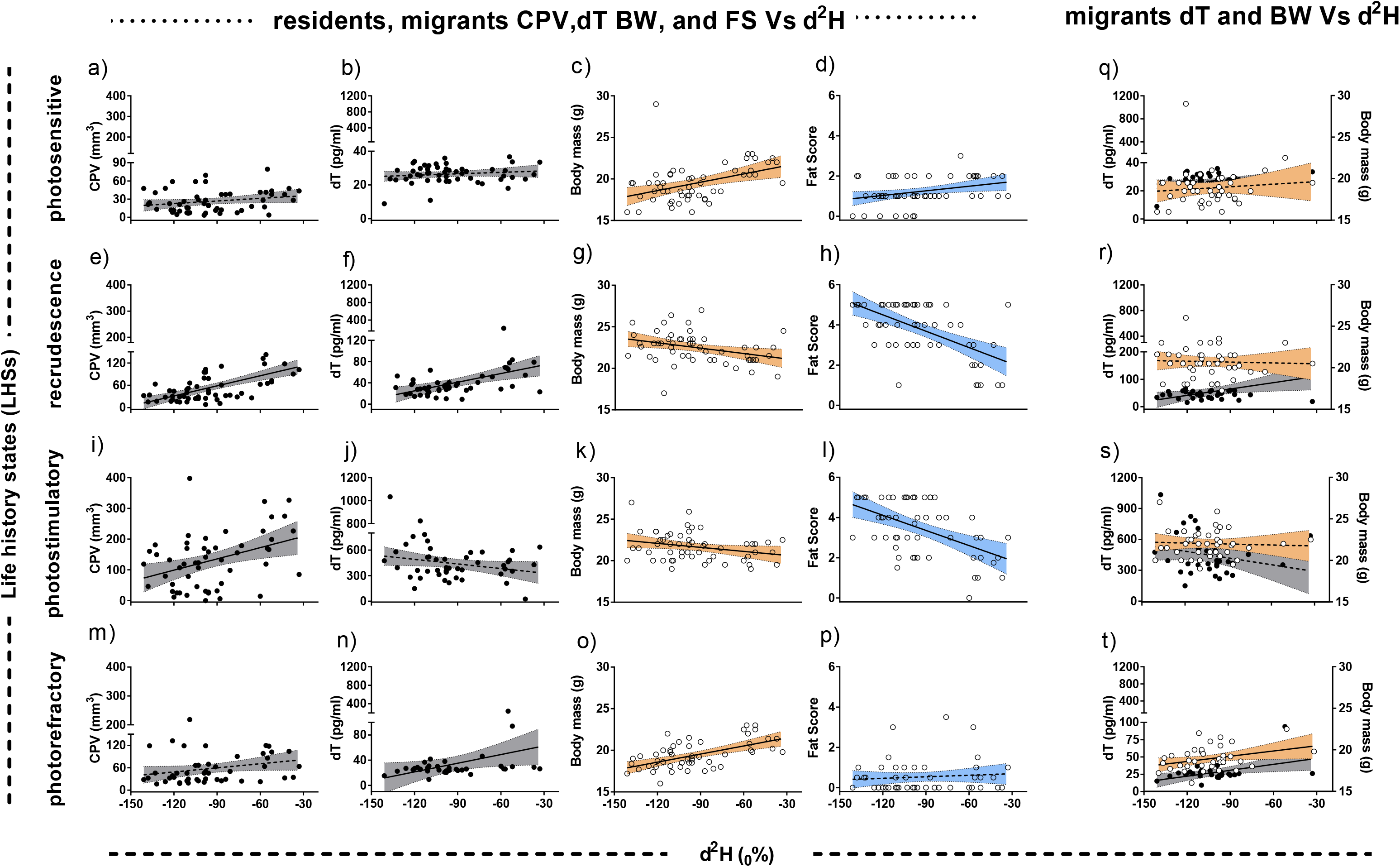
Life history state dependent changes in the relationship between CPV, BW, FS and dT across latitude. Correlation between residents and migrants CPV, T_0_, dT, BW, and FS against δ^2^H as a constant variable in photosensitive (9L: initial point; a-d), recrudescence (12.4L: CPV/BW/FS and 11L: dT; critical photoperiod threshold, e-h), photostimulatory (15L: CPV, dT level at highest peak; i-l), and photorefractory (16L: endpoint; m-p) states. Correlation among migrants dT and BW against δ^2^H values across different LHSs (q-t). Pearson correlation was used for all physiological data except fat score which is an ordinal value. Spearman correlation was used for fat score ordinal data. X-axis represents the δ^2^H as a constant variable and Y axis represents individual data points for CPV, dT (hollow circles), BW, FS (solid circle). Statistical significance was defined by alpha <0.05. Linear regression line with 95% confidence interval (shaded area) represents significant correlation as solid line and dotted line for no significant correlation.

#### Migrants, residents, and stable isotope

During the photosensitive state, δ^2^H values were significantly positively correlated with BW (r = 0.4184, p = 0.0013; Fig. 3c) and FS (r = 0.2695, p = 0.045; Fig. 3 d), but not with CPV or dT (Fig. 3 a, b). During recrudescence, both CPV (r = 0.647, p < 0.0001; Fig. 3 e) and dT (r = 0.4698, p = 0.0006; Fig. 3 f) showed significant positive correlation and BW (r = −0.3315, p = 0.0126; Fig. 3 g) and FS (r = −0.4752, p = 0.0002; Fig. 3 h) showed negative correlation with δ^2^H values at their respective change point day lengths.

When resident juncos reached their peak (the stimulatory phase), δ^2^H values were positively correlated with CPV (r = 0.3823, p = 0.0043; Fig. 3 i) and negatively correlated with BW (r = −0.273, p = 0.0458; Fig. 3 k), and FS (r = −0.4786, p < 0.0001; Fig. 3 l). However, when all the birds reached peak stimulation at 15L, δ^2^H values were not correlated with dT (Fig. 3 j). When the birds reached the refractory state, BW (r = 0.5657, p < 0.0001; Fig. 3 o) and dT (r = 0.334, p = 0.0403; Fig. 3 n) were positively correlated with δ^2^H values providing evidence for latitudinal variation in timing of reaching refractoriness. CPV and FS were no longer correlated with δ^2^H values (Fig. 3 m, p).

#### Migrants and stable isotopes

Migrants with different δ^2^H values did not show any correlation with dT and BW in the photosensitive state (Fig. 3 q). With respect to dT in migrants at 11.75 L (recrudescence; CPP for LLM), gonadal recrudescence was delayed in migrants with lighter δ^2^H values (r = 0.3708, p = 0.0201) and there was no correlation between δ^2^H values and BW (Fig. 3 r). When the migrants reached peak photostimulation, there was no correlation between δ^2^H values and dT, BW (Fig. 3 s). Under the refractory state, comparison of migrant dT (r = 0.4452, p = 0.0155) and BW (r = 0.3264, p = 0.0426) (Fig. 3 t) reveled a positive correlation with δ^2^H values, providing evidence for δ^2^H-dependent difference in the timing of onset of refractoriness which occurred earlier with lower isotope values (i.e., proxy for higher latitude).

### Timing of molt, migrants and residents

Finally, post-breeding molt score was negatively correlated with δ^2^H values during the refractory state (Primary molt: r = −0.4318, p = 0.0008, Fig. 4 a, b; head molt: r = −0.3975, p = 0.006; Fig. 4a-d). Relationship between dT levels, BW and molt score support that birds breeding at different latitudes also differ in the timing of refractoriness (Fig. 4; Fig. 5). Migrants did not show any significant correlation with δ^2^H values during the refractory state.

**Figure 4.**
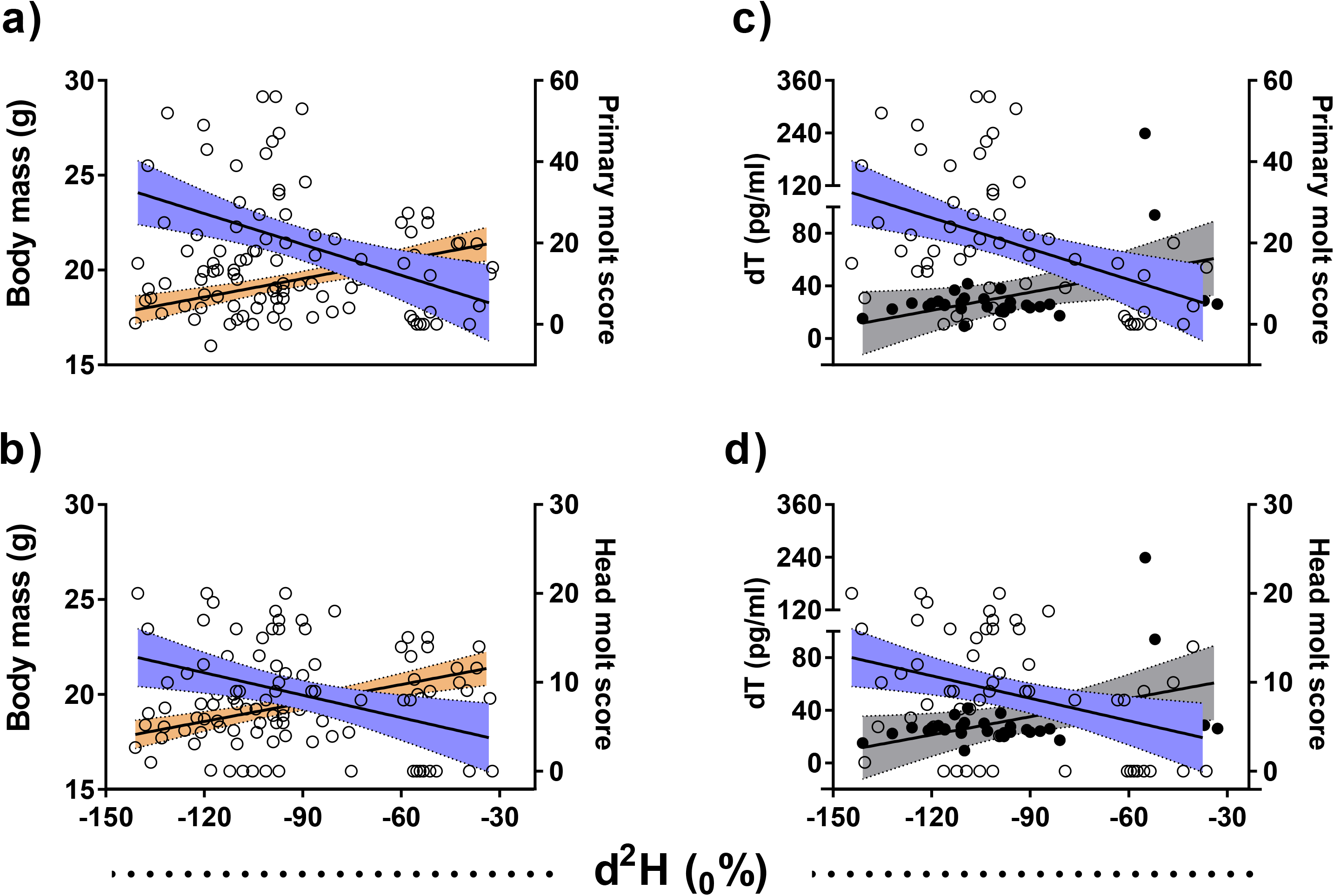
Latitudinal variation in timing of molt, dT, and BW during endpoint refractory state. Pearson correlation between BW/ primary molt score (a), BW/ head molt score (b), dT/primary molt score (c), and dT/head molt score (d) against δ^2^H as a constant variable in 16 L photorefractory state. X-axis represents the δ^2^H as a constant variable and Y axis represents individual data points BW, dT (hollow circles), primary and head molt score (solid circle). Statistical significance was defined by alpha <0.05. Linear regression line with 95% confidence interval (shaded area) represents significant correlation as solid line and dotted line for no significant correlation.

**Figure 5.**
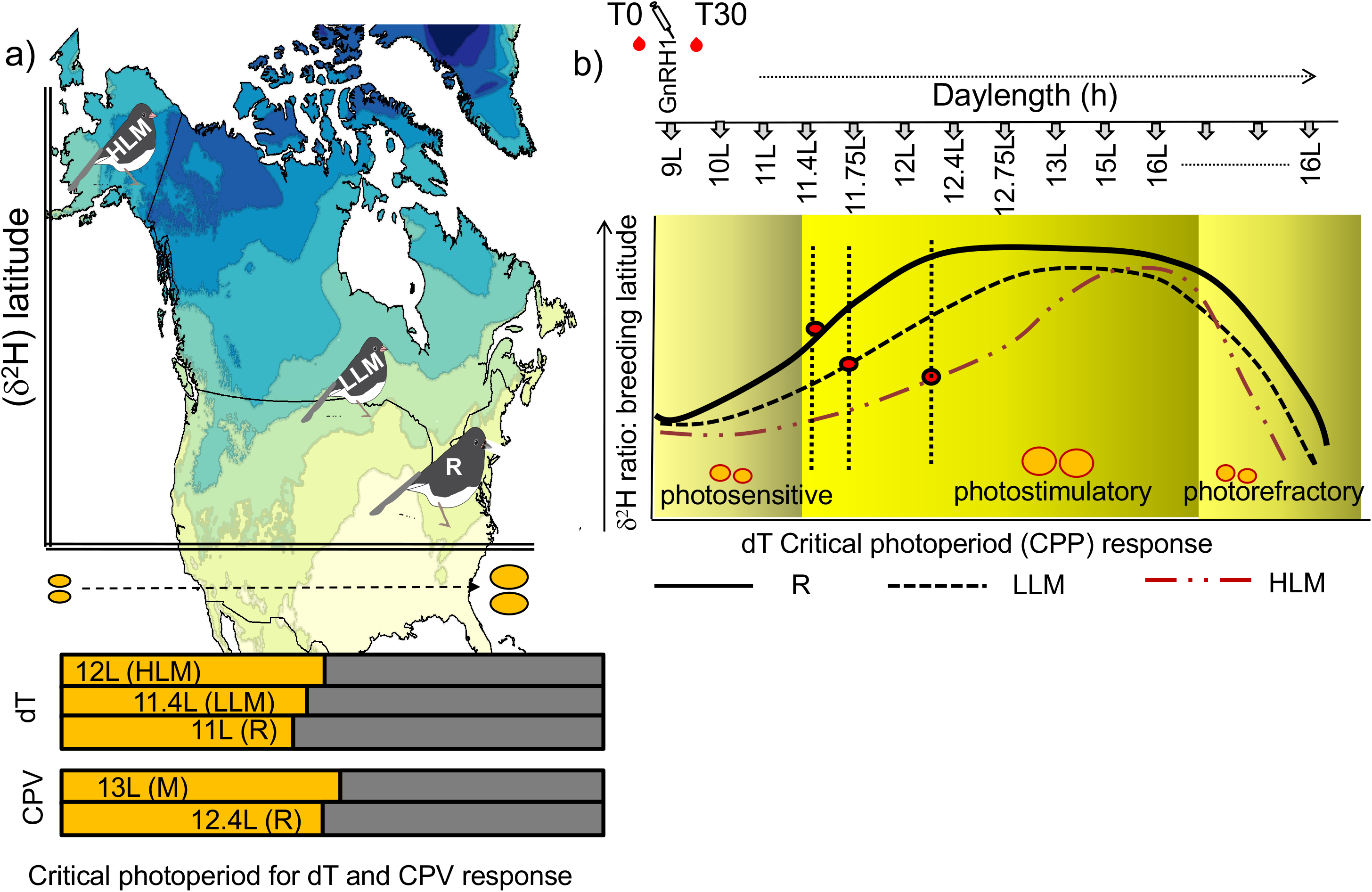
Latitudinal cline in critical photoperiod (CPP) for gonadal recrudescence and pace of life history states. Schematic summary Figure showing the breeding latitude dependent response to different critical photoperiod (CPP) and difference in the pace of different life history states in dark eyed juncos. Left panel showing the CPP for cloacal protuberance and dT response across latitude (a). Right panel showing the latitude-dependent seasonal waveforms of different life history states derived from common garden experiment. Photosensitive juncos overwintering at the same latitude exposed to increasing day length in a common garden show divergence in the development of recrudescence, photostimulation and timing of attaining photorefractoriness. Residents (black solid line), Low Latitude Migrants (black dotted line), high latitude migrants (red dotted line) exhibit differences in CPP to initiate gonad recrudescence, reach stimulation and attain photorefractoriness. Residents breeding at lower latitudes reproduce earlier, maintain stimulatory phase for a longer period and undergo refractory state later. High and Low Latitude migrants delay recrudescence and reach peak stimulation later and undergo refractoriness sooner (b).

## Discussion

We found that captive migrant and resident juncos held under naturally increasing photoperiod exhibited different critical photoperiods for gonadal recrudescence. Resident juncos exhibited earlier CPV growth and elevation in dT than migrants. Further, timing of elevated CPV and dT response were associated with latitude as estimated using stable δ^2^H values. Our findings thus provide evidence for population level variation in the timing of initiation and termination of breeding in a common environment as a function of latitude, when latitude is estimated using δ^2^H isotope. Termination of reproduction started earlier in migrants than residents, as indicated by earlier timing of post-breeding molt, concurrent with earlier decline in the dT levels. Assuming that birds return to their sites of origin, we can conclude that birds destined to breed at higher latitudes delayed onset of gonadal growth as a correlate of breeding later in the year after completing longer migrations. To our knowledge such continuous covariation between the timing of annual life-history states, photosensitive - recrudescence – stimulation, and refractoriness across different breeding latitudes has not been described previously in any bird species (Fudickar et al., 2016; Ramenofsky et al., 2017).

### Birds breeding at lower latitudes initiate preparation for reproduction earlier

The natural history of reproductive timing has shown that birds species breeding at higher latitudes tend to breed later and terminate reproduction sooner. Additionally, within species, populations breeding at higher latitudes initiate egg laying later than closely-related populations found at lower latitudes (Baker 1938, Myers, 1955). Two independent studies investigating seasonal reproductive physiology in free living quail from two different locations differing by 9° latitude showed a 2-3 week advance in egg laying date in birds breeding at lower latitudes (Genelly 1955; Anthony 1970). Another study from two independent labs in England (52°N) and California (37°N) showed maximum stimulated gonads earlier at lower latitudes corresponding to first egg laying dates (Dawson and Goldsmith 1982; Rothery et al., 2001).

With respect to migratory species, previous studies have related migratory distance and breeding latitude to reproductive timing in different bird species (Rubenstein et al., 2002, Fudickar et al., 2016), and numerous studies have shown that within species complexes, populations living at lower latitudes tend to have longer breeding seasons than those found at higher latitude (Dawson and Goldsmith 1983; Dawson, 2013; Greives et al., 2016). But preparation for breeding begins prior to the beginning of breeding season as defined by first egg laying date. Gonadal recrudescence precedes reproduction and is associated with a rise in circulating testosterone.

### Latitude is directly proportional to critical photoperiod response for gonad recrudescence

There are a very few studies testing the difference in photoperiodic threshold as a prerequisite to initiate early gonadal recrudescence. Birds collected from three different latitudes (45°, 57°, 70° N) and maintained in a common garden set up where they were exposed to gradually increasing photoperiod showed earlier maturation in those from the lower latitude (45°N), but not in other two groups of birds from higher latitudes (Silverin et al., 1993). A recent common garden study of two subspecies of white crowned sparrows (*Zonotichia leucophrys*), and dark-eyed juncos (*Junco hyemalis*) that differed in migratory strategy showed differential responses in sensitivity to increasing day length, which influences the induction and termination of breeding (Fudickar et al., 2016; Ramenofsky et al., 2017). Our results fill several gaps in knowledge about the differences in the critical photoperiod threshold of dark-eyed junco subspecies as they transition from photosensitivity to gonadal recrudescence, duration of the stimulatory phase and timing of attaining photorefractoriness.

### Birds breeding at higher latitude terminate reproduction sooner

The transition from the stimulatory to refractory state is signaled by molt (Hall and Fransson 2000) and our results also revealed a difference between migrant and resident juncos in the timing of refractoriness. This compares to starlings held in captivity and exposed to photoperiods simulating annual cycles at higher (52°N) and lower (9°N) latitude. The starlings showed earlier gonadal maturation in 9°N, but birds from both latitudes regressed their gonads at the same time (Dawson, 2013). In our study, towards the end of the experiment when all birds were on 16h photoperiod, high latitude migrants no longer responded to GnRH by elevating T, while residents and low latitude migrants continued to elevate T in response to GnRH. Further, the relationship between molt score and T in response to GnRH across latitude also showed that birds originating from higher latitudes became photorefractory earlier.

The observed difference in the pattern created by latitudinal variation in CPP and seasonal life-history states can provide a framework for testing how various climatic variables account for variation in seasonal timing of birds from different latitudes. The juncos with lighter δ^2^H delayed recrudescence, remained in breeding phase for a shorter period, and become refractory sooner, unlike the juncos with heavier δ^2^H which recrudesced earlier, had longer breeding periods and entered the refractory state later. In total our results point out towards latitudinal variation in the pace of life-history states and the mechanisms underlying seasonal changes in the responsiveness of HPG axis.

Seasonal life-history states and associated phenology have long been studied, but are currently receiving renewed attention in the context of global climate change. In the temperate zone, photoperiod, temperature, and other environmental variables often correlate with seasonal phenology across latitude. As, a consequence, the study of species-specific seasonal phenology has strong application in the context of global climate change (Schwartz 2003, Parmesan 2006). Fitness for a seasonal animal involves not only the ability to synchronize behavior and physiology to the seasons but also to anticipate, prepare, and cope with the changing seasons. For migratory birds, there is an additional challenge of identifying the optimal time to initiate migration and recrudesce gonads while living at locations distant from their breeding locations. For example, warmer winters are resulting in earlier springs. As a consequence, migrants that arrive at times that were formerly optimal find that peak food availability for rearing offspring has already passed, leading to mismatches between migration schedules and optimal times to breed (Lack 1968; Visser at al., 2004). Some long-distance migratory bird species have advanced their spring arrival dates in response to climate change, but others have not and arrived too late for the pulse of insect food needed to nurture offspring (Visser et al., 2004; Jonzén et al., 2006). Mismatch in the seasonal timing has significant consequences at the population level (Nussey et al., 2005; Both et al, 2006). Thus, knowing the mechanisms of life-history state-dependent phenology may help in predicting the effect of climate change on survival and fitness of species (Miller-Rushing et al., 2010).

In order to estimate the ecological consequences of climate change we must be able to forecast the shifts in the direction, magnitude, and phase of phenological processes under different environmental scenarios. This forecasting is impeded due to a shortage of precise knowledge of the mechanisms determining the pace of life-history events. Phenology is a key process that reflects an organism’s micro-evolutionary response to a wide range of environmental cues (van Asch et al., 2007). Animals distributed geographically across a wide range of photoperiod, temperature, and other environmental conditions that vary with latitude are known to express phonological events at different phases of the annual cycle. Hence, it is critical to incorporate mechanistic and evolutionary perspective while forecasting ecological consequences of climate change.

## Conclusions

Our study demonstrates differences in seasonal timing across latitudes in response to changing photoperiod and reveals some of the underlying mechanisms and their potential for adaptive response to environmental change. Birds breeding at lower latitudes recrudesced earlier, maintained the stimulatory phase for longer, and attained refractoriness later. Whereas, birds originating at higher latitude delayed recrudescence, remained stimulatory for a shorter time, and attained refractoriness sooner. That is, latitude was directly proportional to the critical photoperiod required for recrudesce and inversely proportional to the timing of refractoriness. Particularly informative was a group of migrants that had δ_2_H similar to residents and an intermediate response in CPP and timing of refractoriness. This may indicate that migration delays reproduction, but the extend of the delay depends on how far a bird has to travel. The approach used in this study can be applied to other species in which populations that differ in where they breed are found in the same winter environment as they do or do not prepare to migrate. It will be interesting to learn whether their patterns resemble those seen in the junco.

## Declaration

Authors have no conflict of interest.

## Authors’ contribution

DS and EDK conceived the idea. DS, SRR, AAK, and KAA carried out the experiment. CS performed the hydrogen stable isotope experiment and provided data. DS wrote the manuscript with the help of all authors. All authors approved the final draft.

## Acknowledgements

The funds were provided by the Indiana University through the Grand Challenge Initiative, Prepared for Environmental Change to EDK. We thank Adam M. Fudickar for his suggestions during experimental design. We thank Cody Ross Philips and Zhenyue Tan from Indiana Statistics Consulting Center, IU Bloomington for statistical help. We thank Jesse Montoure, Izzy Krahling, Katie M. Talbott and Allison Byrd for helping in blood samplings and molt scoring. We thank Nathan E. Fletcher for assistance with animal care during experiment. We thank David Sinkiewicz for providing access to the Center for Integrative Study of Animal Behavior (CISAB) lab facility.

